# Macrophage microRNA-146a is a central regulator of the foreign body response to biomaterial implants

**DOI:** 10.1101/2024.04.03.588018

**Authors:** Manisha Mahanty, Bidisha Dutta, Wenquan Ou, Xiaoping Zhu, Jonathan S Bromberg, Xiaoming He, Shaik O. Rahaman

## Abstract

Host recognition and immune-mediated foreign body response (FBR) to biomaterials can adversely affect the functionality of implanted materials. To identify key targets underlying the generation of FBR, here we perform analysis of microRNAs (miR) and mRNAs responses to implanted biomaterials. We found that (a) miR-146a levels inversely affect macrophage accumulation, foreign body giant cell (FBGC) formation, and fibrosis in a murine implant model; (b) macrophage-derived miR-146a is a crucial regulator of the FBR and FBGC formation, as confirmed by global and cell-specific knockout of miR-146a; (c) miR-146a modulates genes related to inflammation, fibrosis, and mechanosensing; (d) miR-146a modulates tissue stiffness near the implant during FBR; and (e) miR-146a is linked to F-actin production and cellular traction force induction, which are vital for FBGC formation. These novel findings suggest that targeting macrophage miR-146a could be a selective strategy to inhibit FBR, potentially improving the biocompatibility of biomaterials.

## INTRODUCTION

Biomaterials and medical devices are routinely used in tens of millions of procedures each year as prostheses in orthopedic, dental, cardiovascular, and reconstructive surgery, as scaffold for controlled drug release devices, and as matrix for cell-based therapeutics that is the next pillar of modern medicine (1–6). However, the implantation of biomaterials/devices into soft host tissue often leads to the development of foreign body response (FBR), a chronic inflammatory condition that is characterized by an inner layer of adherent macrophages and/or destructive foreign body giant cells (FBGCs) with an outside layer of fibrotic scar tissue (2–9). FBR ultimately leads to structural or functional implant failure, and may cause harm to, or death of, the patient. FBR poses a vexing medical challenge because there are no therapeutic options that can mitigate the FBR today (2–5, 7–9). Improved understanding of the molecular mechanisms underlying the generation of FBR is the most important step for the development of novel and effective therapeutic strategies that eliminate or reduce the FBR.

Emerging data support a role for stiffness (modulus) of the extracellular and intracellular matrix in numerous cellular processes including gene expression, inflammation, and cell differentiation (10–19). Recent work has shown that the size, shape, and stiffness of implanted materials have a profound impact on the development of FBR (8, 10, 14, 20–23). For example, large spherical hydrogels show a greater reduction of fibrosis (a critical event in FBR) than do small spherical hydrogels (8). Also, increasing hydrogel stiffness augments the collagen capsule thickness surrounding the implant (14). Interestingly, it has been shown that the FBR to PEG-phosphorylcholine hydrogels changes with respect to both the stiffness and zwitterionic content (21). In this regard, our previous work showed that biomaterial-induced FBR is dependent on TRPV4, a mechanosensitive channel of the transient receptor potential vanilloid (TRPV) family (19). Our recent work demonstrated that the development of implant stiffness-induced FBR in vivo and increase of implant-adhered tissue stiffness are reliant on TRPV4, although the precise mechanisms by which TRPV4 modulates FBR remain unknown (10).

Macrophages are recognized as the key orchestrators of the FBR due to their ability to secrete inflammatory proteins, undergo fusion to form FBGCs, synthesize and remodel matrix components, promote the differentiation of the myofibroblasts that remodel the extracellular matrix (ECM), and form the fibrous capsule around the implant (2, 5, 10, 19, 24–28). Thus, it is essential to identify the inter- and intracellular signaling pathways that mediate macrophage-biomaterial interactions and macrophage responses to biomaterials. In this regard, previous reports by our group and others showed that various macrophage functions, including differentiation, FBGC formation, inflammatory marker expression, motility, and phagocytosis, are sensitive to matrix/substrate stiffness, implying that changes in implant and/or matrix stiffness may play a crucial role in the FBR (10, 14, 27–33). Thus, it is important to identify the cellular stiffness sensor of macrophage and the nature of the implant/tissue stiffness-induced inflammatory and fibrotic signals that mediate the link between stiffness and FBR.

MicroRNAs (miRs) are evolutionary conserved, small (∼22-nucleotide), single-stranded, non-coding RNAs that determine the functional fate of coding genes (34). Their emergence as important regulators in virtually all cells/tissues and in numerous cellular processes, including gene expression, inflammation, and cell differentiation, has prompted us to determine their significance in FBR (35–40). Although there are no reports on miR regulation of FBR to biomaterials so far, miRs have been linked to regulation of macrophages in inflammatory settings and cell fusion scenarios, suggesting that miRs may play a pivotal role in regulating FBR (39, 40). In this report, our results show a novel mechanistic link between miR-146a and FBR. We found that miR-146a modulates the FBR to biomaterials by regulating macrophage activation and fibrogenesis in a manner dependent on implant-induced change in tissue stiffness and cellular traction force.

## METHODS

### Reagent and antibodies

We procured primary antibodies for CD68 (Bio-Rad) and α-SMA (Sigma). Secondary antibodies from Jackson Immunoresearch, including goat, rabbit, and mouse antibodies, were also obtained. Further supplies included ProLong Diamond 4′,6-diamidino-2-phenylindole (DAPI), Alexa Fluor 488– and Alexa Fluor 594–conjugated immunoglobulin G (IgG), and Alexa Fluor 594 Phalloidin. Cell culture essentials like Dulbecco’s modified Eagle’s medium (DMEM), fetal bovine serum (FBS), antibiotic-antimycotic, and related reagents were purchased from Gibco. Mouse-recombinant IL-4, GMCSF, and MCSF were sourced from R&D Systems. We acquired mixed cellulose esters (0.45-μm pore size filters, 12 mm) from Millipore. Polyacrylamide (PA)-hydrogels with stiffness levels of 1 kPa and 50 kPa, along with collagen-coated 25 kPa PA hydrogels containing 0.2 μm fluorescent beads, were obtained from Matrigen.

Sodium alginate, purchased from Sigma (St. Louis, MO), underwent purification through sequential chloroform washing and treatment with 1% (w/v) charcoal. Subsequently, it underwent freeze-drying to remove water content. Collagen type I was procured from Corning (NY). For microfluidic device connections, we utilized masterflex transfer tubing, including the MFLX 95802-00 tubing and MFLX06417-11 tubing, both purchased from VWR (Radnor, PA). Additionally, we ordered 1 ml and 10 ml slip tip syringes from Becton, Dickinson (BD, Franklin Lakes, NJ). SU-8 (SU-8 2050) and SU-8 developer were purchased from Kayaku Advanced Materials (Westborough, MA). Furthermore, silicon wafers were sourced from University Wafer (South Boston, MA).

### Animal

We obtained WT C57BL/6 mice from Charles River Laboratories in the USA. The global miR-146a knockout mice (miR-146a^-/-^) were purchased from The Jackson Laboratory in the USA. Additionally, we acquired macrophage-specific miR-146a KO mice from The Jackson Laboratory. For the custom breeding project, The Jackson Laboratory conducted a cryorecovery process utilizing cryopreserved sperm from Mir146tm1Mtm/Mmjax (MMRRC Stock# 36799-JAX) male mice and oocytes from C57BL/6J (Stock# 000664) female mice. The resulting offspring were crossed with B6.129S4-Gt (ROSA) 26Sortm2 (FLP*) Sor/J (Stock# 012930) mice. Subsequently, these mice were bred with B6.129P2-Lyz2tm1 (cre) Ifo/J (Stock# 004781) mice to generate homozygous (Mirs146) hemizygous (LysM-cre) mice, designated as miR-146a^Mac/null^, and homozygous (Mirs146) wild type (LysM-cre) mice, referred to as miR-146a^fl/fl^ (or miR-146a^flox/flox^). All mice were housed in a controlled environment with regulated humidity, temperature, and specific pathogen-free conditions. All experimental procedures and animal usage were conducted in accordance with the approvals granted by the Institutional Animal Care and Use Committee and the University of Maryland Animal Review Committee.

### Generation of FBGC *in vitro* and F-actin staining

Bone marrow was extracted from the femurs of WT, miR-146a^-/-^, miR-146a^fl/fl^, and miR-146a^Mac/null^ mice. The isolated bone marrow cells were cultured in DMEM supplemented with 10% FBS and MCSF (25 ng/ml). Incubation was carried out at 37°C with 5% CO2 for a period of 7-8 days to allow the generation of mature BMDMs (Bone Marrow-Derived Macrophages). For the formation of FBGCs on Permanox slides, a cell density of 500,000 BMDMs was seeded per well in DMEM supplemented with 10% FBS (10, 19). This culture was maintained for 8-10 days, with the additional application of IL4 and GMCSF (25 ng/ml) every 48 hours. To confirm the successful generation of FBGCs, the slides were fixed using a 10% formalin solution, followed by a 4-minute Giemsa stain. To create FBGCs with varying matrix stiffness, 1 x 10^6^ mature BMDMs were seeded onto collagen-coated PA hydrogels with stiffness levels of 1 kPa and 50 kPa (10, 19). This was done in DMEM supplemented with 10% FBS and involved an 8-10 day culture period, with the same IL4 and GMCSF treatment regimen every 48 hours. The hydrogels were fixed using 4% paraformaldehyde, and subsequent immunofluorescence staining was carried out as described below. Additionally, the PA hydrogels were fixed using 4% paraformaldehyde and stained with Alexa-Fluor-labeled phalloidin 594 for F-actin and DAPI for nuclei. Fluorescent imaging was performed at 10x magnification using a Zeiss Axio Observer Z1 microscope.

### In vivo FBR model, histology, and immunofluorescence staining

To induce the formation of FBR in vivo, we performed subcutaneous implantation of either mixed cellulose esters or collagen-coated PA stiffness gels with two different stiffness levels (1 kPa and 50 kPa) (10, 19). After 28 days of implantation, we surgically removed the implants along with the surrounding skin tissue and embedded them in OCT (Optimal Cutting Temperature) compound. These skin tissue samples were then utilized to create tissue sections with a thickness of 7-10 µm using a Cryostat, and these sections were mounted on positively charged slides. Subsequently, the skin tissue sections were fixed using a 10% formalin solution and stained with Masson’s trichrome stain following the manufacturer’s instructions. Additionally, some sections were subjected to staining with anti-CD68 IgG (1:100) or α-SMA (1:100) antibodies at 4°C overnight to detect the presence of macrophages and myofibroblasts in the tissue, respectively. For secondary antibody staining, anti-rabbit or anti-mouse IgG conjugated with Alexa Fluor 488 and Alexa Fluor 594 (1:200) were utilized. DAPI was employed to stain the cell nuclei.

### Adhesion and spreading

Mature BMDMs were placed on coverslips and collagen-coated PA hydrogels with stiffness levels of 1 kPa and 50 kPa. This culture was maintained in DMEM supplemented with 10% FBS. Subsequently, the cells were subjected to treatment, either with or without IL4 plus GMCSF (25 ng/ml), for a duration of 24 hours before being examined under a microscope.

### Reverse transcription quantitative polymerase chain reaction (qRT-PCR)

We isolated total RNA from BMDMs that had been treated with or without IL4 plus GMCSF (25 ng/ml), following the instructions provided in the Qiagen kit. Subsequently, a qRT-PCR was conducted using the One-Step Quantitative Reverse Transcription PCR kit from Bio-Rad. The qRT-PCR utilized specific primers for the following genes: Mcp-1 (qMmuCED0003785), TRAF6 (qMmuCED0046143), TRPV4 (qMmuCID009008), Il1β (qMmuCEP0054181), Il6 (qMmuCED0045760), Tnfα (qMmuCED0004141), CD36 (qMmuCID0014852), TGFβ1 (qMmuCED0044726), and GAPDH (qMmuCEP0039581). To quantify the gene expression levels, we employed the comparative CT method as outlined in the Bio-Rad qRT-PCR user manual. This allowed us to determine the expression of each gene relative to the mRNA levels of GAPDH.

### RNAseq

We began by extracting total RNA from the cellulose ester implant tissue sample, followed by an enrichment step using oligo(dT) magnetic beads to remove rRNA. This enrichment was carried out using the Arraystar rRNA removal kit. Subsequently, we prepared RNAseq libraries using an Illumina kit. After completing the library preparation, we assessed the quality of the libraries using the Agilent 2100 Bioanalyzer. For RNAseq, we utilized the Illumina NovaSeq 6000 platform, and the RNAseq was outsourced to Arraystar in Maryland, USA. To identify differentially expressed transcripts (DET) and differentially expressed genes (DEG), we applied criteria of a *p*-value < 0.05, a log2FoldChange of less than −0.585, and a log2FoldChange of more than 0.585. The DET data was employed to create heatmaps using the heat mapper web tool at http://heatmapper.ca/. Additionally, we conducted GO function analysis, KEGG pathway analysis, and KEGG pathway enrichment analysis using the integrated differential expression and pathway analysis web tool (iDEP ver. 0.96) and the pathview tools (41–44).

### Traction force microscopy

BMDMs were placed onto collagen-coated 25 kPa PA hydrogels containing 0.2 μm fluorescent beads in 24-well plates. The cells were allowed to incubate overnight at 37°C with 5% CO2. Subsequently, the cells were subjected to treatment with IL4 and GMCSF (25 ng/ml) for 48 hours. Using a microscope with a 20x objective, fluorescent images were captured both before and after trypsinization, which was performed using a 0.6% SDS solution. To quantify the displacement and traction force generated, we utilized the TractionForAll® software (45).

### Atomic force microscopy

We employed the JPK Nano Wizard 4 atomic force microscope (AFM) from Bruker Nano GmbH in Berlin, Germany, to measure the stiffness of skin tissues adjacent to implants with stiffness levels of 1 kPa and 50 kPa in miR146a^fl/fl^ and miR146a^Mac/null^ mice (10). For this purpose, we utilized 7 μm-thick skin sections immobilized on glass slides submerged in PBS to record individual force spectroscopy curves (F-D curves) in contact mode for each sample. To perform these measurements, we used CP-qp-CONT-BSG-B-5 colloidal probes (sQube), which had specific characteristics, including a nominal resonance frequency of 30 kHz in air, a spring constant of 0.1 N/m, gold coating on the detector side, and an attached borosilicate glass sphere with a diameter of 10 μm. We set a relative set point of 1 nN, an extension speed of 10 μm/s, and a Z length of 10 μm to ensure appropriate F-D curves. A total of 50 F-D curves were recorded across various areas surrounding the implants on each tissue sample. Subsequently, these curves were processed using the Hertz contact mechanics model for spherical probes to determine the Young’s modulus (E). For data visualization and analysis, we used Graph Pad Prism to generate histograms, bar plots, and line plots for each condition.

### Fabrication of microfluidic device

The fabrication of non-planar microfluidic devices involved a soft lithography process using polydimethylsiloxane (PDMS) on a mold or master, which itself was created using photolithography techniques as previously referenced (46–51). The design of the microfluidic device was generated using AutoCAD (Autodesk, Mill Valley, CA), and a corresponding design mask was produced by CAD/Art Service (Brandon, OR). The mold preparation comprised several key steps. Initially, a 100 µm thick layer of SU-8 2050 photoresist was applied via spin-coating onto a 4-inch silicon wafer. The coated wafer underwent heating on a hot plate at temperatures of 65 and 95°C. Subsequently, ultraviolet (UV) light exposure was performed through the patterned mask using a MA-4 Mask Aligner (Karl Suss, Munich, Germany). Following post-exposure baking, an additional 50 µm thick layer of SU-8 2050 was applied, followed by baking at 65 and 95°C and exposure to UV light to create the shell channel pattern. Another 50 µm thick layer of SU-8 2050 was applied and subjected to similar baking and UV exposure to pattern the oil channel. Finally, the wafer was developed using SU8 developer and underwent post-baking at 120°C. The specific parameters for spin speeds, baking, and exposure energy were determined based on the SU-8 2050 datasheet supplied by the manufacturer.

PDMS was prepared by mixing the base-to-curing agent at a ratio of 10:1 (w/w). Both the upper and lower parts of the microfluidic device were created by pouring PDMS over the mold, followed by degassing and baking at 75°C for 2 hours. The PDMS with the designed pattern was then carefully removed from the mold, plasma-treated using a PDC-32G plasma cleaner (Harrick Plasma, Ithaca, NY) at 18 W and 27 Pa for 2 mins, and aligned under a Zeiss Primovert microscope (Oberkochen, Germany). Subsequently, the devices underwent a 3-hour bake at 75°C before being utilized further.

### Microencapsulation of BMDMs in 3D alginate-collagen microcapsules and generation of FBGCs

The core channel fluid was prepared by combining 200 µl of neutralized collagen type I (6 mg/ml) with 200 µl of a 4% sodium alginate solution (46–51). This mixture was used to dissociate BMDMs to achieve the desired concentration, and the resulting mixture was then transferred to a 1 ml syringe. In contrast, the shell fluid consisted of a 2% sodium alginate solution. For the oil channel fluid, we prepared an emulsion by combining mineral oil and a 1 g/ml aqueous calcium chloride solution (volume ratio: 5 to 1, supplemented with 1.5% Span 80). This emulsion was created through sonication for 1 min using a Branson 450 digital sonifier (Emerson, St. Louis, MO). All the fluids were introduced into the microfluidic device via a syringe pump (Harvard Apparatus, Holliston, MA). The flow rates were set at 200 µl/h for the core channel, 500 µl/h for the shell channel, and 5 ml/h for the oil channel. The microcapsules were collected in a 50-ml tube containing a 0.25 M D-mannitol solution (with 10 mM HEPES, pH 7.4). After removing the mineral oil, the microcapsules underwent a wash with a 0.25 M D-mannitol solution, followed by gelation using a 200 mM calcium CaCl2 solution for 10 minutes. They were then washed two more times with a 0.25 M D-mannitol solution before being resuspended in completed DMEM. This medium contained 10% FBS to support further culture in a humidified incubator set at 37°C with 5% CO2. To generate microcapsules with varying core materials or cell numbers, the same procedures as mentioned above were followed, except for the preparation of the core fluid. Instead of mixing 200 µl of neutralized collagen type I (6 mg/ml) with 200 µl of a 4% sodium alginate solution, 200 µl of neutralized collagen type I (6 mg/ml) was mixed with 200 µl of a 1% sodium alginate solution to resuspend different cell counts.

Mature BMDMs were encapsulated within sodium alginate beads with varying core materials and cell numbers to produce FBGCs using the IL4 plus GMCSF (25 ng/ml) fusogenic cocktail every 48 hours for 10 days. The sodium alginate beads were fixed using 8% paraformaldehyde and then stained with Alexa-Fluor-labeled phalloidin 594 and DAPI for visualizing F-actin and cell nuclei, respectively. Fluorescent imaging was performed at 20x magnification using a PE spinning disk confocal microscope.

### Live/dead staining

After various incubation durations, we gathered the alginate microcapsules and subjected them to staining with Calcein AM (Corning, Inc., Corning, NY) and Propidium iodide (Invitrogen, Carlsbad, CA) for a 15-min period. Following this staining process, the samples were washed three times with DMEM. Subsequently, we analyzed the samples using a fluorescence microscope (Zeiss LSM 710) with a 10x phase lens.

### Spinning disk confocal microscopy

To produce fluorescent 3D images of the FBGCs, we employed a PerkinElmer spinning-disk confocal microscope at 20x magnification. The data acquisition was carried out using the PerkinElmer Velocity software (version 6.4.0), which was equipped with a Hamamatsu ImagEM X2 EM-CCD camera (C9100-23B). Images were captured utilizing 405 nm (DAPI) and 561 nm (F-actin) wavelength lasers, with a Z-slice interval of 1 µm.

### Statistical analysis

All data are presented as mean ± SEM. Statistical comparisons between different groups were conducted using Student’s t-test or one-way ANOVA followed by the Bonferroni test. In the notation, “ns” indicates not significant, * denotes p < 0.05, ** denotes p < 0.01, and *** denotes p < 0.001. It’s worth noting that all experiments involving animals, both in vivo and in vitro, were carried out in a randomized and blinded manner.

## RESULTS

### MiR-146a levels inversely affect macrophage accumulation, FBGC formation, and collagen generation in subcutaneous implantation model, with global miR-146a deletion in mice exacerbating these effects

To investigate the global mechanisms involved in FBR to biomaterials, we analyzed microRNA (miR) expression profiles in skin tissue RNA from mice. This included both sham (non-implanted, mock surgery) and mice implanted with cellulose ester filters (0.45 μm pore size, 12 mm^2^) using RNAseq (10, 19, 52). The study utilized a subcutaneous (s.c) implant model in C57BL/6 mice, employing cellulose ester as a common model implant (10, 19, 52). Applying stringent criteria of at least a 1.5-fold change and a significance level of *p* ≤ 0.05, we identified 78 miRs as upregulated and 172 miRs as downregulated in the implanted group compared to the sham group (Fig. 1, A and B). We focused on miR-146a, which was markedly downregulated (∼10-fold) in the implanted mice (Fig. 1, B and C), as it is known to be a potent negative regulator of inflammatory and fibrotic responses (53–56). We observed that decreased miR-146a expression in tissues adhering to the implant correlated inversely with increased collagen accumulation, FBGC formation, and macrophage accumulation (Fig. 1, C, D, E, and F).

**Figure 1.**
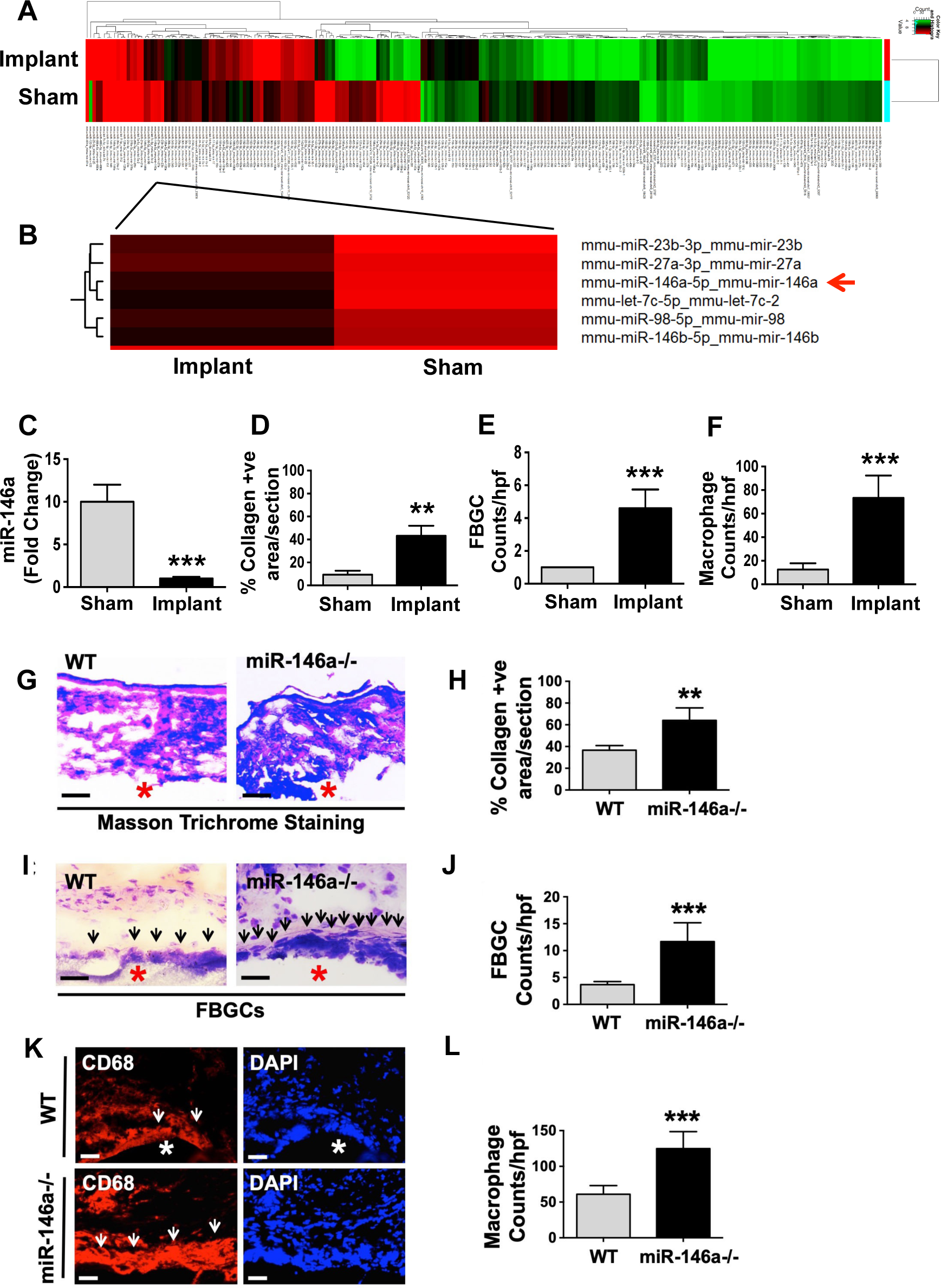
miR-146a expression levels inversely correlates with macrophage accumulation, FBGC formation, and collagen generation in subcutaneous implantation model, with global miR-146a deficiency in mice worsening these effects. Total RNA was extracted from the skin tissues of both sham-operated and cellulose ester-implanted wild-type (WT) mice, 14 days after implantation. **(A, B)** We conducted RNA-Seq to profile miRNA expression. **(C)** qRT-PCR analysis was then carried out to evaluate miR-146a levels, with normalization against sno202 small nucleolar RNA (snRNA) levels. The FBR in the skin tissue of mice was examined 28 days post-implantation. **(D)** Tissue sections underwent Masson’s trichrome staining to quantify collagen deposition. **(E)** FBGCs were identified and quantified using Hematoxylin and Eosin **(H & E)** staining. **(F)** The accumulation of macrophages in the skin sections was quantified after staining with CD68 IgG. Representative images depict sections of filters (marked by asterisks) that were subcutaneously implanted in WT and miR-146a^-/-^ mice for 28 days. **(G)** These sections were also stained with Masson’s trichrome. **(H)** Collagen deposition was quantitatively analyzed (blue staining in G). **(I)** FBGCs (indicated by arrows) were stained using H & E. **(J)** Quantitative data on the number of FBGCs was compiled. **(K)** Sections stained with CD68 IgG (red) and DAPI (blue) highlight macrophages at the tissue-implant interface (arrows). **(L)** The accumulation of macrophages was quantitatively assessed. For each group, n = 5 mice were used. Statistical significance was determined using Student’s t test, with **p ≤ 0.01 and ***p ≤ 0.001. The term ‘hpf’ refers to high power field.

The role of miR-146a in FBR in vivo was further examined using a subcutaneous biomaterial implant model in global miR-146a knockout (miR-146a^-/-^) mice compared to wild-type (WT) mice, with cellulose ester used as the implant material (10, 19, 52). After 28 days, miR-146a^-/-^ mice exhibited a 2-fold increase in collagen deposition (Fig. 1, G and H), a 4-fold increase in the number of FBGCs (Fig. 1, I and J), and a 3-fold increase in macrophage accumulation at the tissue-implant interface (Fig. 1, K and L), compared to WT mice. These findings suggest that the absence of miR-146a exacerbates biomaterial-induced FBR in vivo.

### MiR-146a deletion in macrophages augments macrophage accumulation, FBGC formation, myofibroblast accumulation, and collagen generation in subcutaneous implantation model

To examine the specific role of macrophage-derived miR-146a in FBR, we generated two mouse models: miR-146a^fl/fl^ (control) and miR-146a^Mac/null^ mice. The miR-146a^Mac/null^ mice were created by crossbreeding miR-146a^fl/fl^ mice with Lyz2-Cre strains, as depicted in supplementary Fig. 1A. We validated the macrophage-specific deletion of miR-146a through quantitative RT-PCR analysis of RNA extracted from peritoneal macrophages. This analysis revealed overexpression of the miR-146a target gene, Traf6, in macrophages from miR-146a^Mac/null^ mice (supplementary Fig. 1B). These novel macrophage-specific knockout mice are invaluable for investigating the distinct contributions of macrophage miR-146a in FBR and FBGC formation in vivo.

We assessed the impact of macrophage miR-146a on FBR using the subcutaneous implant model, comparing miR-146a^Mac/null^ mice to miR-146a^fl/fl^ mice with cellulose ester implants (10, 19, 52). After 28 days, the miR-146a^Mac/null^ mice exhibited a 2-fold increase in collagen deposition (Fig. 2, A and B), a 2.5-fold increase in the number of FBGCs (Fig. 2, C and D), a 3-fold increase in macrophage accumulation at the tissue-implant interface (Fig. 2, E and F), and a 2-fold increase in myofibroblast accumulation (Fig. 2, G and H), compared to miR-146a^fl/fl^ mice. These findings indicate that the absence of miR-146a in macrophages intensifies the biomaterial-induced FBR in vivo.

**Figure 2.**
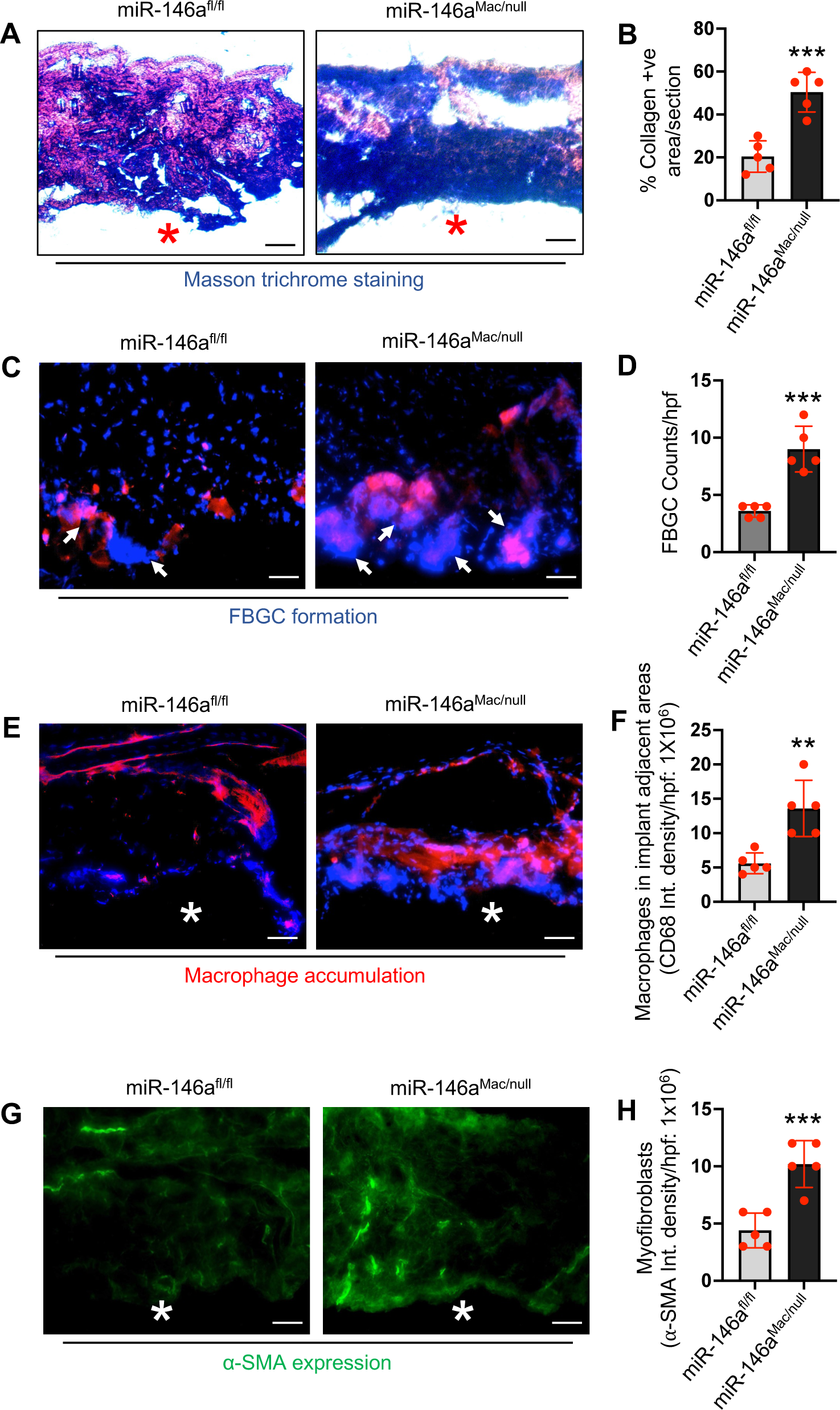
miR-146a deletion in macrophages enhances macrophage accumulation, FBGC formation, myofibroblast accumulation, and collagen production in subcutaneous implantation model. The FBR was evaluated in skin tissue from miR-146a^fl/fl^ and miR-146a^Mac/null^ mice, 28 days after implantation. (**A**) Tissue sections underwent Masson’s trichrome staining, highlighting the collagen production. (**B**) The quantity of collagen produced was measured; red asterisks denote the tissue-implant interface. (**C**) For macrophage visualization, sections were stained with CD68 IgG (red), and nuclei were counterstained with DAPI (blue). Arrows point to macrophages located at the tissue-implant interface. (**D**) This was followed by a quantitative analysis of FBGCs. (**E**) Additional sections were stained with CD68 IgG (red) for macrophage visualization, with nuclei counterstained using DAPI (blue). White asterisks indicate the tissue-implant interface, highlighting macrophage accumulation. (**F**) The findings from E were quantified. (**G**) To identify myofibroblasts, sections were stained with the myofibroblast marker α-SMA (green), and white asterisks mark the tissue-implant interface. (**H**) Quantitative analysis of myofibroblast presence, as indicated in G. For each experimental group, n = 5 mice were used. Statistical analysis was conducted using Student’s t test, with significance levels marked as **p ≤ 0.01 and ***p ≤ 0.001.

### Matrix stiffness-induced foreign body response is reliant on macrophage miR-146a

To evaluate the influence of macrophage-derived miR-146a on matrix stiffness-induced FBR in vivo, we employed a subcutaneous biomaterial implantation model. This model involved implanting collagen-coated polyacrylamide (PA) gel discs with varying stiffness into miR-146a^Mac/null^ mice and miR-146a^fl/fl^ (control) mice. The selected PA discs had either a soft (1 kPa) or rigid (50 kPa) stiffness to replicate the stiffness of normal skin or breast tissue (∼1 kPa) and fibrotic skin (∼8-50 kPa), as published previously (10, 14, 57, 58).

Four weeks post-implantation, we observed a notable difference in collagen deposition around the implants. In miR-146a^fl/fl^ mice, the collagen deposition was 4 times greater surrounding the 50 kPa matrix compared to the 1 kPa matrix. This suggests that fibrosis generation, a key aspect of FBR, is exacerbated by increased implant stiffness (Fig. 3, A and B). Conversely, in miR-146a^Mac/null^ mice, there was a 2.5-fold increase in collagen deposition with the 50 kPa matrix compared to the 1 kPa matrix. Notably, this increase was 1.6 times greater than that observed in miR-146a^fl/fl^ mice with 50 kPa implants. These results imply that the development of stiffness-induced fibrosis in vivo is significantly amplified in the absence of miR-146a (Fig. 3, A and B).

**Figure 3.**
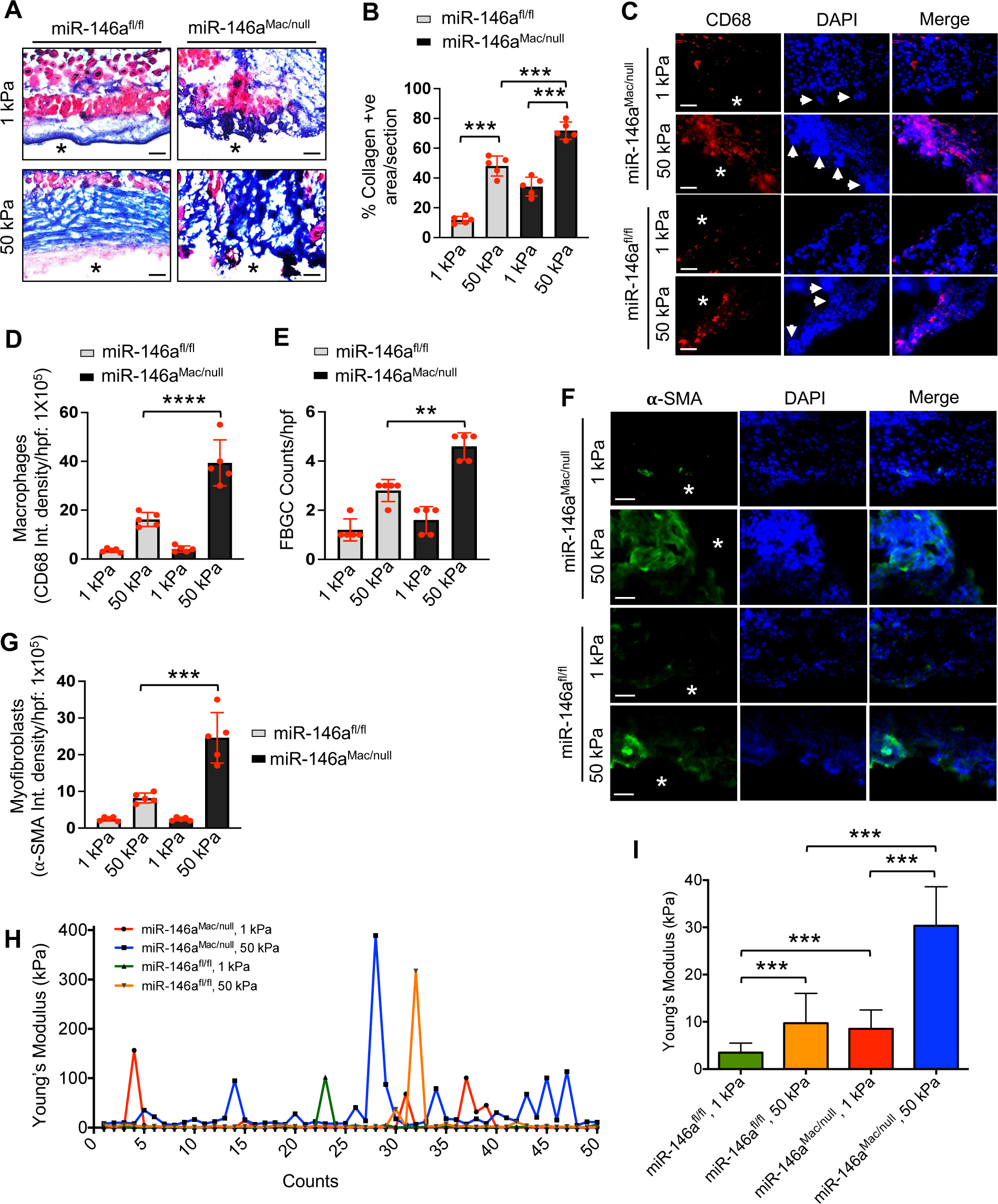
Matrix stiffness-induced FBR depends on macrophage miR-146a. (**A**) Images of representative skin sections from miR-146a^fl/fl^ and miR-146a^Mac/null^ mice implanted subcutaneously with 1 kPa and 50 kPa collagen-coated PA gels (asterisks indicated tissue-implant interface) after 28 days. Sections were stained with Masson’s trichrome to examine the deposition of collagen (blue color). (**B**) Quantification of collagen positive areas in skin sections from the experiment shown in A. (**C**) Skin tissue sections from miR-146a^fl/fl^ and miR-146a^Mac/null^ mice with 1 kPa and 50 kPa implants were stained with anti-CD68 IgG to show macrophage accumulation and multinucleated FBGCs (white arrow) near the tissue-implant interface (white asterisks). Nuclei were stained with DAPI (blue). (**D, E**) Quantification of macrophage accumulation and FBGC formation at the tissue-implant interface in skin sections from the experiment shown in C. (**F**) Skin tissue sections from miR-146a^fl/fl^ and miR-146a^Mac/null^ mice with 1 kPa and 50 kPa implants were stained with anti-α-SMA IgG to show myofibroblast accumulation. Nuclei were stained with DAPI (blue). (**G**) Quantification of myofibroblast accumulation in skin sections from the experiment shown in F. n = 5 mice/group; 2 implantation/mice; Scale bars: 50 μm for (A, C and F); Student’s *t*-test; \*\**p* < 0.01, \*\*\**p* < 0.001, \*\*\*\**p* < 0.0001. (**H**) Plot profile of Young’s modulus of the dataset generated by AFM analysis. n = 5 tissue samples/group and 50 force curves/sample. **(I**) Quantification of Young’s modulus of the dataset generated by AFM analysis of the skin tissues from miR-146a^fl/fl^ and miR-146a^Mac/null^ mice with 1 kPa and 50 kPa implants. n = 5 tissue samples/group and 50 force curves/sample. One-way ANOVA followed by Bonferroni test; \*\*\**p* < 0.001.

The implantation of biomaterials into soft tissues initiates the recruitment and accumulation of inflammatory cells, particularly macrophages, around the implant surface and adjacent areas (10, 19). These cells often merge to form FBGCs, which can damage the implant. In our study, both macrophage accumulation and FBGC formation on the surfaces of 50 kPa implants were significantly increased in miR-146a^fl/fl^ mice, by 8-fold and 3-fold respectively, compared to 1 kPa implants (Fig. 3, C, D, and E). In contrast, miR-146a^Mac/null^ mice showed even more pronounced changes: a 15-fold increase in macrophage accumulation and a 2.5-fold increase in FBGC formation with 50 kPa implants compared to 1 kPa. These increases were 2.6 times (for macrophage accumulation) and 1.6 times (for FBGC formation) higher than those observed in miR-146a^fl/fl^ mice with 50 kPa implants. This indicates that the absence of miR-146a in macrophages significantly amplifies stiffness-induced macrophage accumulation and FBGC formation in vivo (Fig. 3, C, D, and E). These findings suggest that the severity of stiffness-induced FBR in vivo is markedly increased without macrophage miR-146a. Furthermore, the observation of reduced collagen deposition, diminished macrophage accumulation, and fewer FBGCs in miR-146a^fl/fl^ mice carrying soft implants, along with the exacerbation of these effects in miR-146a^Mac/null^ mice, implies that the response to biomaterial stiffness in FBR is miR-146a-dependent in macrophages.

In order to investigate the involvement of fibroblasts/myofibroblasts in generating tissue stiffness, we conducted experiments using a subcutaneous implant model. This model compared miR-146a^Mac/null^ mice to miR-146a^fl/fl^ mice, using collagen-coated PA hydrogel discs with stiffness levels of 1 kPa and 50 kPa. Four weeks post-implantation, we observed a marked difference in myofibroblast presence (as indicated by α-SMA expression levels) around the implants. In miR-146a^fl/fl^ mice implanted with the 50 kPa matrix, myofibroblast abundance was 3 times higher compared to those receiving the 1 kPa matrix. This suggests that myofibroblast recruitment and differentiation, which are key aspects of FBR, are intensified by increased implant stiffness (Fig. 3, F and G). Conversely, in miR-146a^Mac/null^ mice, there was a 10-fold increase in myofibroblast abundance with the 50 kPa matrix compared to the 1 kPa matrix. Notably, this increase was 2.5 times greater than that observed in miR-146a^fl/fl^ mice with 50 kPa implants. These findings indicate that the differentiation of myofibroblasts in response to increased stiffness is significantly more pronounced in the absence of macrophage-derived miR-146a (Fig. 3, F and G).

To investigate the relationship between tissue stiffening adjacent to implants and myofibroblast differentiation in vivo in mice, we conducted measurements of fibrous capsule tissue stiffness using atomic force microscopy (AFM). This was performed on skin tissues implanted with PA hydrogel. Twenty-eight days after implantation, we carried out AFM analysis on skin tissue sections surrounding the implants to derive force curves representing tissue stiffness. Our findings revealed that, four weeks post-implantation, the stiffness of the fibrous capsule tissue around the implant was three times higher in miR-146a^fl/fl^ mice that received the 50 kPa matrix compared to those with the 1 kPa matrix. This indicates that the tissue stiffness elicited by the implant is intensified by a stiffer matrix (Fig. 3, H and I). In comparison, miR-146a^Mac/null^ mice implanted with the 50 kPa matrix exhibited a fourfold increase in tissue stiffness compared to those with the 1 kPa implants. Moreover, this increase was 3.5 times greater than the increase observed in miR-146a^fl/fl^ mice with 50 kPa implants. These results suggest that tissue stiffening induced by implant stiffness in vivo is significantly more pronounced in the absence of miR-146a in macrophages (Fig. 3, H and I).

### Macrophage miR-146a differentially modulates expression of inflammatory, fibrotic, and mechanosensitive genes in biomaterial-implanted skin tissues

To investigate the impact of macrophage miR-146a on gene expression related to FBR, we conducted RNAseq and qRT-PCR analyses. Initial sequencing data was quality-checked using FASTQC software, as illustrated in Supplementary Fig. 2, A and B. Here, the y-axis represents the quality score (Q) and the x-axis the position in the sequence read, with the blue line indicating the mean Q score. A score of 30, shown in the green area, signifies good quality. Post quality control, we assessed the protein coding potential using the Coding Potential Assessment Tool (CPAT), with coding and non-coding transcripts represented as red and blue dots, respectively, in the 3D scatter plot (Supplementary Fig. 2C) (2,3). Principal Component Analysis (PCA) was then utilized to simplify the large dataset and align gene expression clustering (Supplementary Fig. 2D). This identified 3275 DEGs with a p-value < 0.05, highlighting variations between miR-146a^Mac/null^ and miR146a^fl/fl^ implant tissues (Supplementary Fig. 2, E and F).

The sequencing results revealed 1951 DETs with a p-value < 0.05 and log2FC±0.585, of which 916 were upregulated and 1035 downregulated in miR-146a^Mac/null^ compared to miR-146a^fl/fl^. The top 200 upregulated and downregulated DETs were visualized using the Heatmapper tool (Fig. 4A). Further, heatmaps for DETs related to fibrosis, inflammation, and mechanosensing were generated (Fig. 4, B, C, and D), accompanied by tables detailing descriptions, p-values, fold changes, and biological functions (Table 1). Notably, we observed increased expression of integrins, Stat1, Mmp13, Bcl2, and Nlrp10 in miR-146a^Mac/null^ compared to miR-146a^fl/fl^. To delve deeper into miR-146a’s role in FBR development, we treated both miR-146a^Mac/null^ and miR-146a^fl/fl^ BMDMs with an IL4 plus GMCSF fusogenic cocktail for 24 hours, and subsequent qRT-PCR analysis revealed a significant increase in genes linked to fibrosis, inflammation, and macrophage fusion, such as Cd36, Il1β, Tgfβ1, Tnfα, and Trpv4 in miR-146a^Mac/null^ (Fig. 4, E-L).

**Figure 4.**
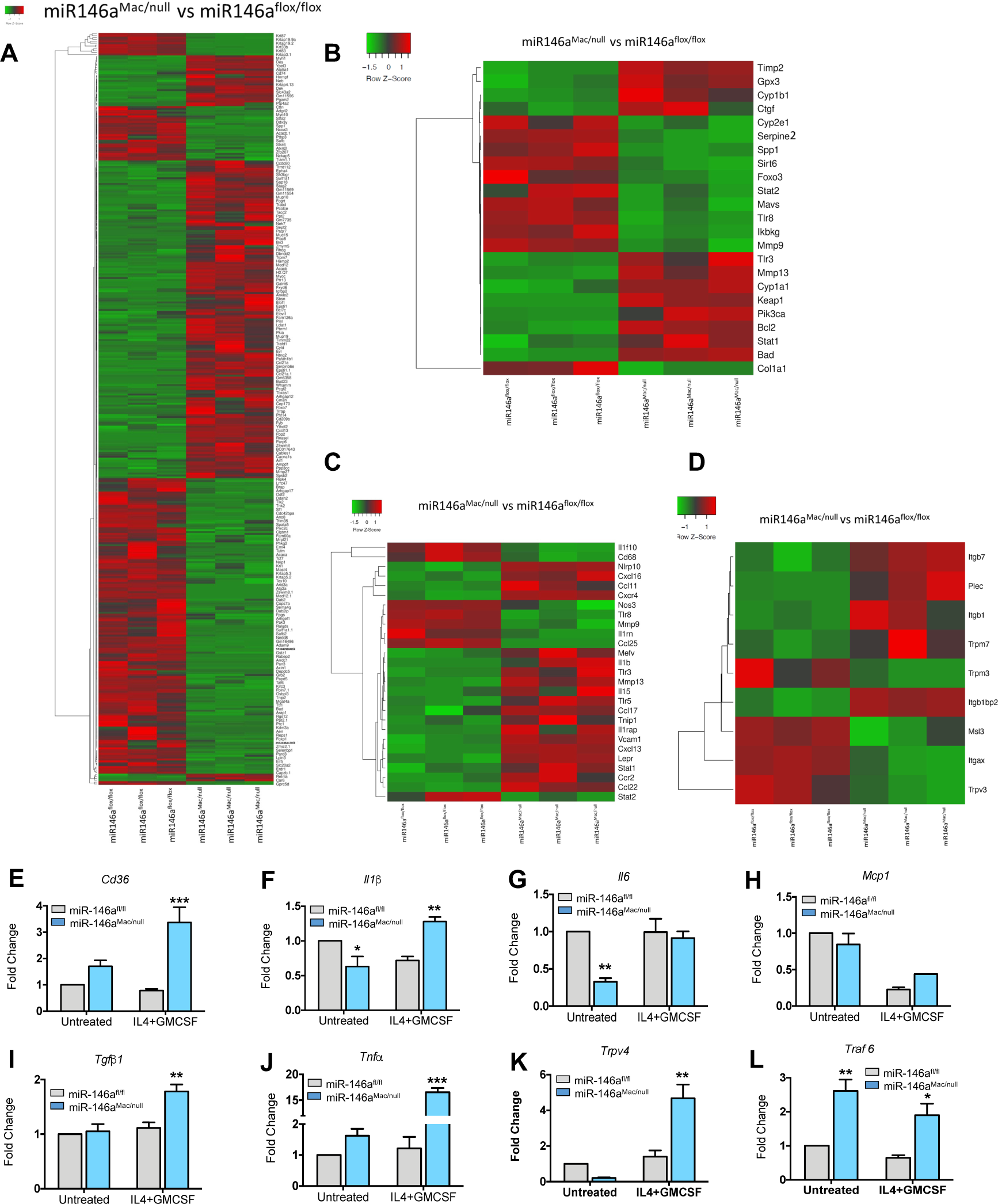
Macrophage miR-146a variably modulates inflammatory, fibrotic, and mechanosensitive gene expression in biomaterial-implanted skin tissues. RNAseq analysis was performed on cellulose ester-implanted skin tissues from miR-146a^fl/fl^ and miR-146a^Mac/null^ mice, 28 days post-implantation. The analysis pooled samples from 5 mice per group. (**A-D**) This analysis focused on: (**A**) A heatmap showcasing the expression patterns of the top 200 most upregulated and 200 most downregulated DETs. (**B-D**) Separate heatmaps depicting the expression patterns of DETs specifically associated with fibrosis, inflammation, and mechanosensitivity. Further, gene expression analysis was conducted on BMDMs from miR-146a^fl/fl^ and miR-146a^Mac/null^ mice, treated with or without IL4 plus GMCSF for 24 hours. This analysis included the genes Cd36 (**E**), Il1β (**F**), Il6 (**G**), Mcp1 (**H**), Tgfβ1 (**I**), Tnfα (**J**), Trpv4 (**K**), and Traf6 (**L**), using qRT-PCR. Each condition was replicated 4 times for statistical robustness. The data were analyzed using one-way ANOVA followed by a Bonferroni post-hoc test, with significance levels noted as *p < 0.05 and **p < 0.01.

**Table 1.**
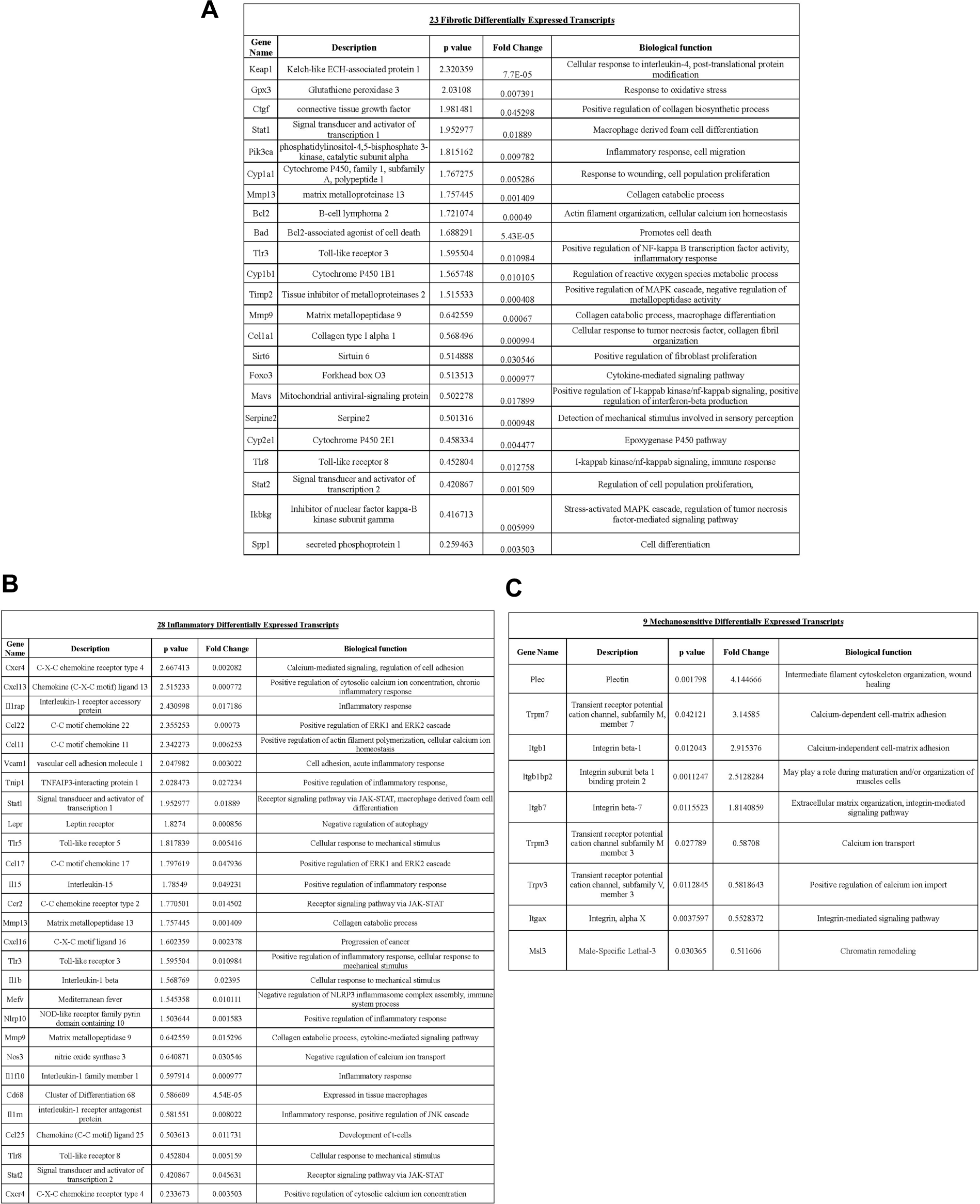
List of selected genes differentially expressed in miR-146a^Mac/null^ as compared to miR-146a^fl/fl^. Table of 23 fibrotic DETs (**A**), 28 Inflammatory DETs (**B**), and 9 mechanosensitive DETs (**C**) with their descriptions, *p*-value, fold change, and biological functions are shown.

Gene Ontology (GO) enrichment analysis of the 3275 DEGs highlighted the top 10 significantly enriched GO terms in each category (biological processes, cellular components, and molecular functions), showcasing a range of cellular responses including immunity, myofibril formation, contractile fiber involvement, mannose binding, and actin binding (Supplementary Fig. 3, A-H). Subsequently, we selected 1951 DETs based on a p-value < 0.05 and log2FC±0.585, mapping them to 2 KEGG pathways. This mapping revealed an upregulation in the gene expression of PFN, MLC, PI3K, ITG, CXCR4, and TGFβRII in miR-146a^Mac/null^ (Supplementary Fig. 4, A and B).

### MiR-146a regulates matrix stiffness-induced macrophage spreading, adhesion, and FBGC formation

Adhesion and spreading, key factors in macrophage accumulation, were markedly enhanced on stiff substrates (50 kPa) in miR-146a^-/-^ BMDMs compared to their WT counterparts, as shown in Fig. 5, A, B, and C. This suggests that the absence of miR-146a function contributes to increased macrophage activation in response to substrate stiffness. We further investigated miR-146a’s role in modulating FBR in BMDMs in vitro, employing an IL4 and GMCSF-induced FBGC formation assay (10, 19). Our findings revealed that miR-146a^-/-^ BMDMs exhibited a three-fold increase in FBGC numbers and a two-fold increase in cell fusion compared to WT (Fig. 5, D, E, and F). Similarly, miR-146a^Mac/null^ BMDMs showed a 1.6-fold increase in FBGC numbers, a two-fold increase in cell fusion, and a 1.7-fold increase in the average size of FBGCs compared to miR-146a^fl/fl^ (Fig. 5, G-J).

**Figure 5.**
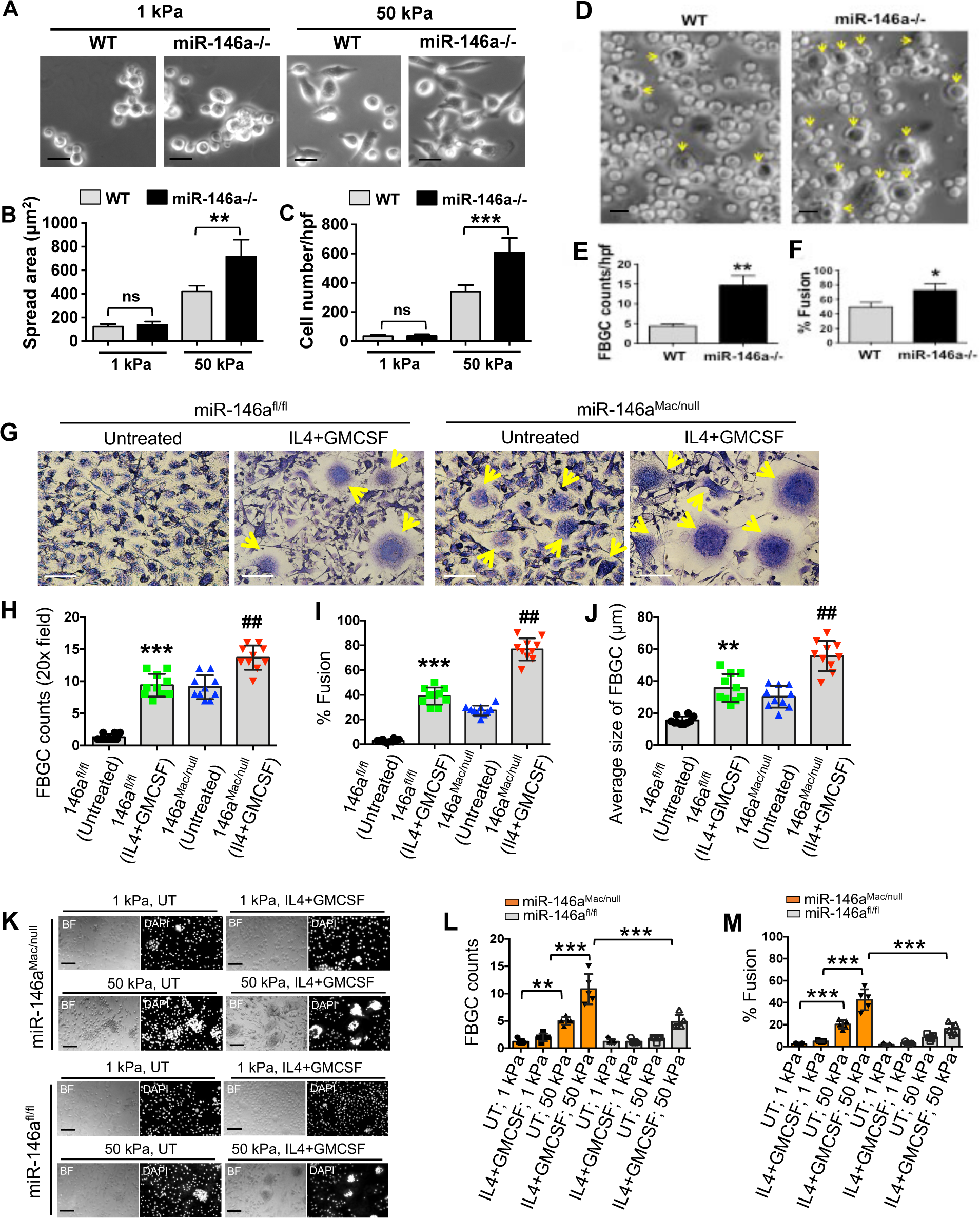
miR-146a modulates matrix stiffness-induced macrophage spreading, adhesion, and FBGC formation. (**A**) Images of cells show increased spread area in miR-146a^-/-^ BMDMs plated on collagen-coated (10 µg/ml) 1 kPa or 50 kPa PA hydrogels for 48h compared to WT. Mean spread area (**B**), and adhered cell number/field (n= 10 fields/condition) (**C**) were quantified (**p ≤ 0.01, ***p ≤ 0.001; Student’s t test; ns: not significant). (**D**) Increased FBGC formation (indicated by arrows) in miR-146a^-/-^ BMDMs compared to WT after IL4 plus GMCSF (25 ng/ml) treatment. Number of FBGC/hpf) (**E**), and % macrophage fusion (**F**) was quantified. \**p* ≤ 0.05, \*\**p* ≤ 0.01, Student’s t test. (**G**) Images of Giemsa-stained multinucleated FBGCs of untreated or cytokine-treated miR-146a^fl/fl^ or miR-146a^Mac/null^ BMDMs plated on Permanox slides. (**H-J**) Quantification of data from the experiment shown in G. (**H**) number of FBGCs, **(I**) % BMDM fusion, and (**J**) the average size of FBGC were shown. n = 3 biological replicates, 6 images/group; scale bars: 50 μm; Student’s *t*-test, **/^##^*p* < 0.01, \*\*\**p* < 0.001. (**K-M**) BMDMs from miR-146a^fl/fl^ and miR-146a^Mac/null^ mice were seeded on collagen-coated (10 µg/ml) 1 kPa and 50 kPa PA gels treated with or without IL4 plus GMCSF for 8 days to generate FBGCs. Representative brightfield and DAPI images (**K**), number of FBGCs (**L**), and % BMDM fusion (**M**) were shown. n = 3 biological replicates, 5 images/group; scale bars: 20 μm, One-way ANOVA followed by Bonferroni test; \*\**p* < 0.01, \*\*\**p* < 0.001.

To confirm if our in vitro findings mirrored in vivo macrophage-related activity, we explored the role of miR-146a in FBR in BMDMs on collagen-coated PA hydrogels with physiological softness (1 kPa) and stiffness (50 kPa), using an IL4 and GMCSF-induced FBGC formation assay (10, 19, 27). Significant upregulation of FBGC formation was observed on stiffer PA gels compared to softer ones in miR-146a^Mac/null^ macrophages (Fig. 5, K, L, and M). IL4 and GMCSF priming further augmented FBGC numbers and fusion percentage (Fig. 5, K, L, and M). These alterations in FBGC formation, influenced by stiffness and soluble factors, were less pronounced in miR-146a^fl/fl^ macrophages, underscoring a pivotal role of miR-146a in stiffness-sensitive FBGC formation under physiologically relevant conditions.

### MiR-146a-dependent F-actin generation and cellular traction force induction are associated with FBGC formation

Cytoskeletal remodeling is central to various physiological processes, including cell fusion. Notably, the generation of cellular traction force is primarily governed by F-actin production, a key aspect of cytoskeletal remodeling (59, 60). To explore if miR-146a-dependent matrix stiffness-induced FBGC formation is linked to cytoskeletal remodeling, we assessed F-actin levels in IL4 plus GMCSF induced FBGCs using a 2D PA hydrogel in vitro model. Our findings revealed a notable increase in F-actin production on stiffer PA gels compared to softer ones in miR-146a^Mac/null^ macrophages (Fig. 6, A and B). IL4 plus GMCSF treatment further amplified F-actin levels (Fig. 6, A and B). However, these stiffness- and soluble factor-related changes in F-actin production were less pronounced in miR-146a^fl/fl^ macrophages (Fig. 6, A and B), indicating miR-146a’s significant role in stiffness-responsive F-actin generation under physiologically relevant conditions.

**Figure 6.**
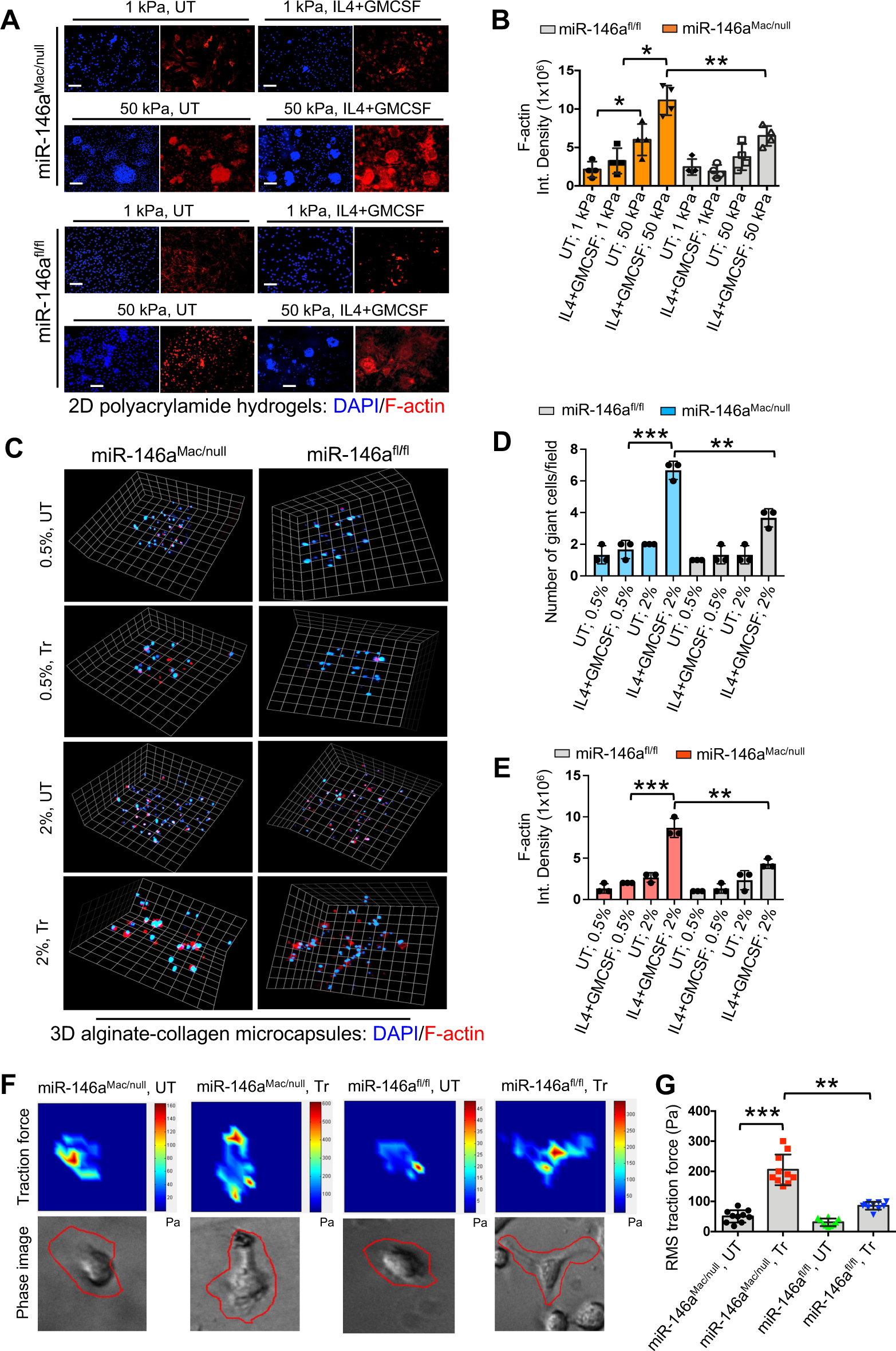
miR-146a-dependent F-actin formation and cellular traction force induction are critical in FBGC formation. (**A**) Images of miR-146a^fl/fl^ and miR-146a^Mac/null^ BMDMs plated on collagen-coated (10 µg/ml) 1 kPa or 50 kPa PA hydrogels with or without 25 ng/ml of IL4 plus GMCSF for 8 days show phalloidin-stained F-actin (red) and DAPI-stained nuclei (blue). (**B**) Quantification of data from the experiment shown in A. n = 3 biological replicates, 5 images/group; scale bars: 50 μm, One-way ANOVA followed by Bonferroni test; \**p* < 0.05, \*\**p* < 0.01. (**C**) Spinning disk confocal microscopy images show a 3D view of miR-146a^fl/fl^ or miR-146a^Mac/null^ BMDMs seeded inside soft (0.5%) or stiff (2%) 3D alginate-collagen microcapsules and cultured with or without 25 ng/ml of IL4 plus GMCSF for 10 days. Images show phalloidin-stained F-actin (red) and DAPI-stained nuclei (blue). Quantification of multinucleated FBGCs numbers (**D**) and F-actin levels (**E**) from the experiment shown in C. (**F**) Color-coded traction vector maps indicate the magnitude of the traction vector and corresponding phase images show the cells. (**G**) Graph shows quantitation of results from traction force microscopy assay of miR-146a^fl/fl^ and miR-146a^Mac/null^ BMDMs plated on collagen-coated PA hydrogels (25 kPa) with or without IL4 plus GMCSF for 24h. 10 cells/group; One-way ANOVA followed by Bonferroni test; \*\**p* < 0.01, \*\*\**p* < 0.001. UT: untreated; and Tr: Treated.

Recent studies have indicated that various physical aspects of implanted materials, such as size, shape, stiffness, and porosity, can influence the intensity of the FBR (8, 10, 14, 20–23). We extended our investigation to assess the impact of implant stiffness on FBGC formation and F-actin production using a 3D alginate-collagen microcapsule in vitro model. Alginate hydrogel, commonly employed in cell and tissue encapsulation for cell-based therapies, was used in this experiment (46, 47, 61–63). We observed a marked increase in both FBGC formation and F-actin production in stiffer 2% alginate microcapsules compared to softer 0.5% alginate capsules in miR-146a^Mac/null^ macrophages (Fig. 6, C, D, and E). IL4 plus GMCSF priming further elevated FBGC and F-actin levels (Fig. 6, C, D, and E). These changes, influenced by stiffness and soluble factors, were more moderate in miR-146a^fl/fl^ macrophages (Fig. 6, C, D, and E), underscoring the crucial role of miR-146a in stiffness-sensitive FBGC formation and F-actin production.

To determine if miR-146a-dependent FBGC formation was associated with cellular traction force generation, we employed traction force microscopy to measure force generation in macrophages induced by IL4 plus GMCSF. We found that IL4 plus GMCSF priming significantly increased traction force generation in miR-146a^Mac/null^ macrophages compared to miR-146a^fl/fl^ macrophages (Fig. 6, F and G), highlighting miR-146a’s essential role in cellular force generation. Collectively, these results suggest that F-actin-dependent cytoskeletal remodeling and the subsequent increase in cellular force may be instrumental in miR-146a-dependent FBGC formation.

## DISCUSSION

The molecular mechanisms driving the FBR remain largely elusive, presenting a significant challenge in medical science. Strategies to mitigate FBR to biomaterials and devices have included various approaches, such as coating devices with basement membrane-derived hydrogels, localized administration of steroids and growth factors, employing alginate-based and zwitterionic material-based hydrogels, and identifying novel molecular targets (20, 64–70). While these methods have shown promise, the results are sometimes inconsistent or contradictory, as documented in various studies (20, 64–70). Advancing these critical technologies, coupled with a more detailed understanding of the molecular basis of FBR, is essential for fundamentally addressing and potentially resolving FBR issues at their root.

While there have not yet been any documented instances of miR involvement in the FBR to biomaterials, there is evidence linking miRs to the regulation of macrophages in contexts of inflammation and cell fusion (39, 40). This suggests a potential significant role for miRs in the regulation of FBR. By identifying and understanding the miRs that are pivotal in the FBR, there is an opportunity to create targeted therapeutic strategies. For example, the use of miR mimics or inhibitors to adjust the levels of certain miRs could effectively modulate the body’s reaction to implanted materials. This approach holds potential for mitigating adverse effects, such as excessive inflammation or fibrosis, associated with the FBR.

Prior to this study, the specific influence of miR expression, particularly miR-146a, on the FBR to biomaterial implants had not been determined. MiR-146a is known as a potent inhibitor of inflammation and fibrosis (53–56), which are key factors in FBR (2–9). Our study addresses this gap in knowledge by showing that mice deficient in miR-146a exhibit an intensified FBR. This finding underscores the critical role of miR-146a in modulating FBR and opens new avenues for therapeutic strategies targeting miR-146a to improve implant success. The specific contribution of macrophage-derived miR-146a to FBR was previously unknown. Our research, using miR-146a^Mac/null^ mice, reveals that the absence of miR-146a in macrophages leads to a significantly increased FBR. This finding is pivotal in identifying cell-specific molecular targets, such as macrophage-derived miR-146a, for enhancing biomaterial compatibility and reducing adverse implant reactions.

Prior research has established that the stiffness of implanted materials significantly influences the FBR, but the detailed mechanisms behind this relationship were not fully understood (8, 10, 14, 20–23). Our study addresses this gap by exploring how miR-146a affects FBR in relation to the varying stiffness of biomaterials. We discovered a direct correlation between the stiffness of biomaterial matrices and the development of fibrosis, with stiffer matrices eliciting more pronounced fibrotic responses. This indicates that the stiffness of biomaterials can significantly amplify fibrotic reactions. Our findings shed new light on the potential role of macrophage miR-146a in cellular responses to material stiffness. Specifically, we observed enhanced fibrosis and myofibroblast differentiation in miR-146a^Mac/null^ mice, which suggests a critical role of macrophage-derived miR-146a in modulating the body’s response to the stiffness of biomaterials. This insight is crucial for biomaterial design, as it implies that controlling implant stiffness might help in modulating FBR for improved biocompatibility.

Our findings not only highlight the impact of matrix stiffness on fibrosis and inflammation but also underscores the complex interactions between fibroblasts/myofibroblasts and biomaterials, contributing to tissue stiffness. The use of atomic force microscopy in measuring tissue stiffness provides a more nuanced understanding of these interactions. This study paves the way for future investigations, particularly in manipulating miR-146a expression to control FBR. Additionally, it calls for further research into the molecular mechanisms by which miR-146a perceives stiffness and influences fibrotic responses, offering new directions for enhancing the success of biomaterial implants.

This study delves into the role of macrophage miR-146a in modulating gene expressions linked to the FBR. RNAseq and qRT-PCR analysis revealed a significant number of DEGs, demonstrating substantial genetic variation between miR-146a^Mac/null^ and miR-146a^fl/fl^ implant tissues. The identification of 3275 DEGs and 1951 DETs marks a critical advancement in understanding the genetic complexities of FBR. The observed upregulation of genes associated with fibrosis, inflammation, and mechanosensing, particularly integrins, Stat1, Mmp13, Bcl2, and Nlrp10 in miR-146a^Mac/null^, provides new insights into the genetic contributors to FBR.

Additionally, the analysis reveals crucial gene ontology terms tied to immunity and myofibril structure and highlights the upregulation of essential pathways such as MLC, PI3K, ITG, CXCR4, TGFβRII in miR-146a^Mac/null^ through KEGG mapping, providing insights into miR-146a’s role in macrophages affecting gene expression related to FBR. The study’s insights into miR-146a’s influence on key genes and pathways significantly enhance our understanding of FBR at the genetic level.

Furthermore, this study illuminates a significant gap in the FBR field by revealing the critical role of miR-146a in macrophage behavior towards substrate stiffness and FBGC formation. The findings that miR-146a deficiency enhances macrophage activation, fusion, and FBGC formation on stiff substrates offer new clarity on the mechanistic underpinnings of FBR. Priming BMDMs with IL4 and GMCSF led to an increase in FBGC numbers, size, and fusion percentage in miR-146a^Mac/null^ BMDMs, indicating that both substrate stiffness and soluble factors collaboratively influence FBGC formation in a miR-146a-dependent manner. Previously, the impact of physical cues on macrophage-mediated FBR was not fully understood. Now, the observed increase in adhesion, spreading, and FBGC formation in miR-146a^-/-^ BMDMs suggests that targeting miR-146a could be a novel strategy for modulating FBR, particularly in the context of medical implants. These insights have profound implications for designing biomaterials that could mitigate adverse FBR by manipulating miR-146a expression or function, presenting a promising direction for future research and therapeutic interventions.

Cytoskeletal remodeling, particularly F-actin generation, is central to various physiological processes including cell fusion (59, 60). Using physiologically relevant 2D PA hydrogel and 3D alginate-collagen microcapsule models, we demonstrated a role of miR-146a in F-actin generation and FBGC formation, in response to substrate stiffness. It reveals that the absence of miR-146a in macrophages leads to enhanced F-actin formation and FBGC generation on stiffer substrates, suggesting miR-146a’s regulatory influence over cell adaptation to mechanical cues. This finding is pivotal as it connects the dots between substrate stiffness, cytoskeletal dynamics, and FBR severity, an area not fully elucidated before. These results align with previous findings that the physical characteristics of implanted materials, such as stiffness, shape, and porosity, critically influence FBR severity (8, 10, 14, 20–23). Moreover, by demonstrating that both substrate stiffness and soluble factors like IL4 and GMCSF amplify F-actin levels, FBGC formation, and cellular force generation in a miR-146a-dependent manner, the study provides new insights into the mechanistic pathways of FBR. These insights are critical for designing biomaterials with optimized stiffness to modulate FBR effectively, offering a new direction for improving biocompatibility and therapeutic outcomes in medical implants.

We cannot exclude the possibility that the reduced FBR and FBGC formation in macrophages in global miR-146a null mice is associated with developmental compensation or unknown secondary compensation. Our studies utilizing miR-146a^Mac/null^ mice mitigated this concern. Four weeks post-implantation, we discovered that implant stiffness affects myofibroblast abundance, an effect mediated by miR-146a, indicating its role in modulating stiffness-induced myofibroblast differentiation during FBR. Atomic force microscopy analysis revealed that tissue stiffness near implants in miR-146a^fl/fl^ mice was tripled with stiff versus soft implants, and in miR-146a^Mac/null^ mice, it quadrupled with stiff versus soft implants, highlighting miR-146a’s significant impact on stiffness-induced tissue stiffening. Macrophages and fibroblasts, both sensitive to matrix stiffness variations (10–15), coexist in inflamed fibrotic tissue (2–9), hinting at their potential interaction influenced by miR-146a-dependent biochemical factors and matrix stiffness during FBR. Given miR-146a’s anti-inflammatory and anti-fibrotic properties in various cells, including macrophages, the regulation of FBR likely extends beyond matrix stiffness sensing, mediated through miR-146a.

Enhancing the host acceptance of implanted biomaterials/medical devices, a key to unlocking various medical breakthroughs, currently hinges on the use of broad-spectrum anti-inflammatory drugs (70–72). These drugs are employed for immune suppression or the management of long-term implantation of biomedical devices (70–72). However, their lack of specificity for a particular immune cell population, coupled with diverse targets and effects in vivo, presents a challenge. This underscores the necessity for the discovery and development of more precise drug targets and their associated inhibitory compounds. Since miRs have unique mechanisms of action, the ability to function as master regulators of the genome, and an apparent lack of toxicity in normal cells/tissues, several miR-based therapeutics are currently under development (73). The results from this study may allow us to uncover miR mechanisms that operate in FBR, which may provide a mechanistic basis to facilitate the translation of this important new knowledge into the clinical arena. Further, since macrophage activation is essential to tissue maintenance and to the development of fibrotic disorders of the liver, heart, lung, skin, and kidney, understanding the role of macrophage miR-146a is likely to lay the foundation for designing novel strategies to aid wound healing and to target miR-146a therapeutically in various diseases including the FBR and fibrosis.

In summary, our study identifies miR-146a as a pivotal regulator of the immune-mediated FBR to implanted biomaterials, influencing macrophage accumulation, giant cell formation, and fibrosis. Targeting miR-146a offers a novel approach to improve biomaterial biocompatibility by mitigating adverse immune responses.

## Acknowledgements

This work was supported by an NIH (R01EB024556) grant to Shaik O. Rahaman. We acknowledge the use of ChatGPT 4 for proofreading the final draft in March 2024.

## Author contributions

SOR, MM, BD, WO, and XH conceived the study, designed and performed the experiments, and analyzed data. SOR and MM wrote the manuscript. SOR, JSB, XH, and XZ edited the manuscript. All authors reviewed and approved the final content of the manuscript.

## Data availability

All data generated and used during this study are included in this article.

---

**Supplementary Fig. 1.**
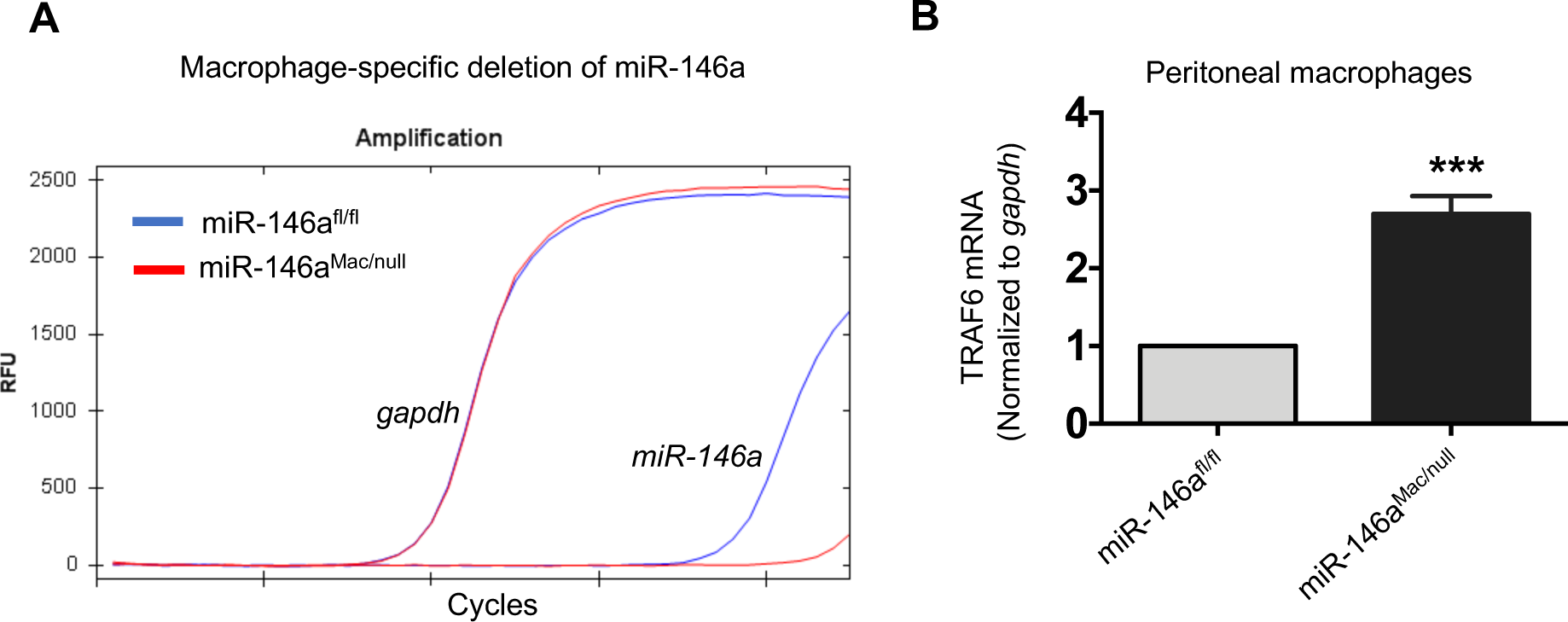
Generation of miR146a^fl/fl^ and miR146a^Mac/null^ mice. **(A)** Confirmation of macrophage-specific deletion of miR-146a by qRT-PCR. **(B)** qRT-PCR analysis shows miR-146a target gene, TRAF6, is overexpressed in peritoneal macrophages from miR146a^Mac/null^ mice.

**Supplementary Fig. 2.**
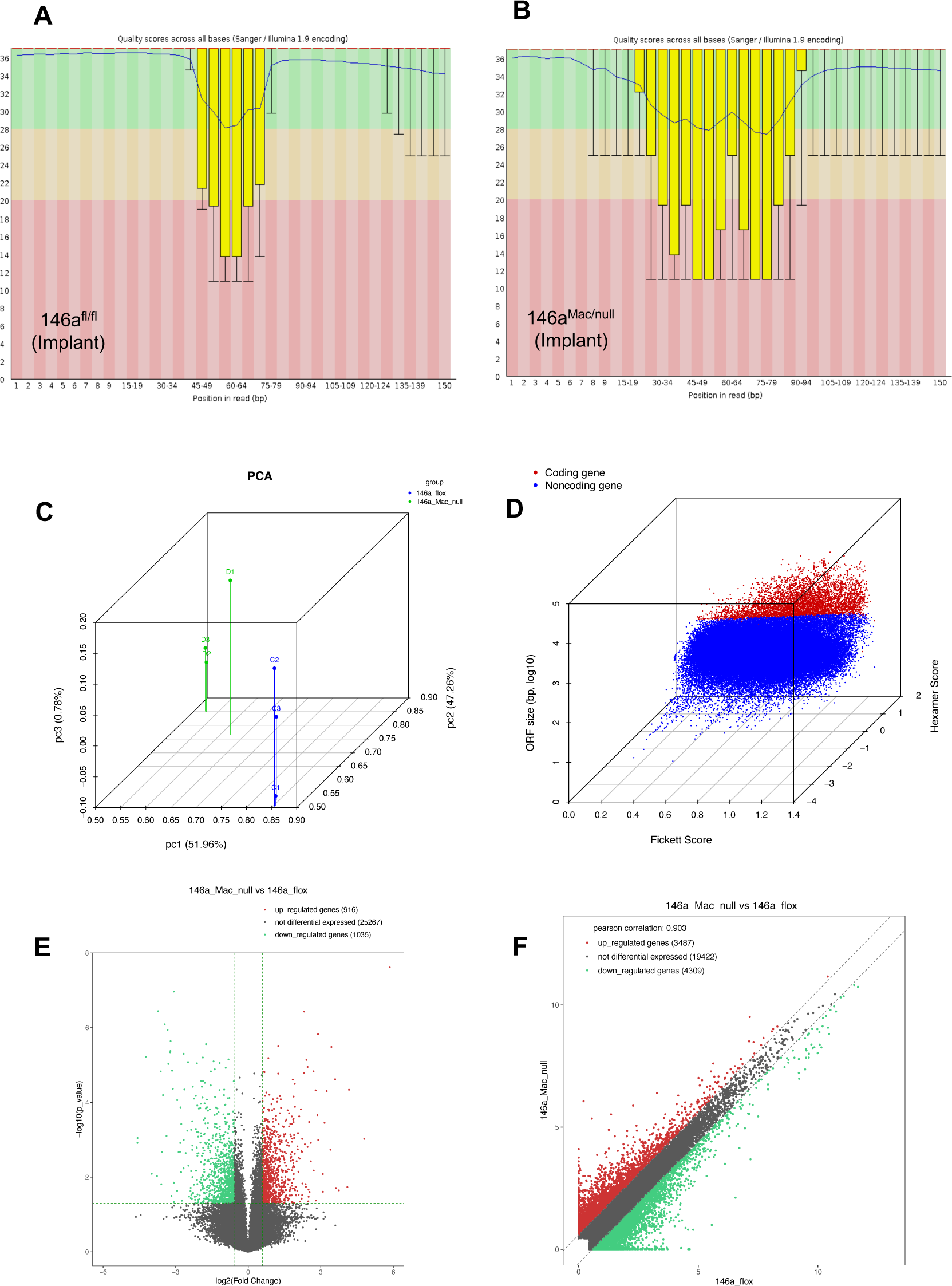
FASTQC software was used to determine the RNAseq quality before downstream processing. RNAseq analysis was conducted on cellulose ester-implanted skin tissues from miR-146a^fl/fl^ and miR-146a^Mac/null^ mice, 28 days after implantation. This analysis involved pooled samples from five mice in each group. The key aspects of the analysis included: (**A, B**) A quality score plot was generated, showcasing the distribution of quality scores (Q) along the y-axis against the read position on the x-axis. (**C**) A 3D scatter plot was constructed to differentiate between coding and non-coding transcripts. This plot utilized the Fickett score, Hexamer score, and open reading frame (ORF) size as parameters, with coding transcripts marked in red and non-coding transcripts in blue. (**D**) Principal component analysis (PCA) was employed to analyze the differentially expressed genes (DEGs). (**E**) A scatter plot of DEGs was created, based on their log2(FPKM) values. (**F**) A volcano plot was generated for the DEGs, plotting log2(fold Change) on the x-axis against −10log10(p-value) on the y-axis. In these plots, the quality score Q is calculated as −10log10(P), where P is the probability of a base call error.

**Supplementary Fig. 3.**
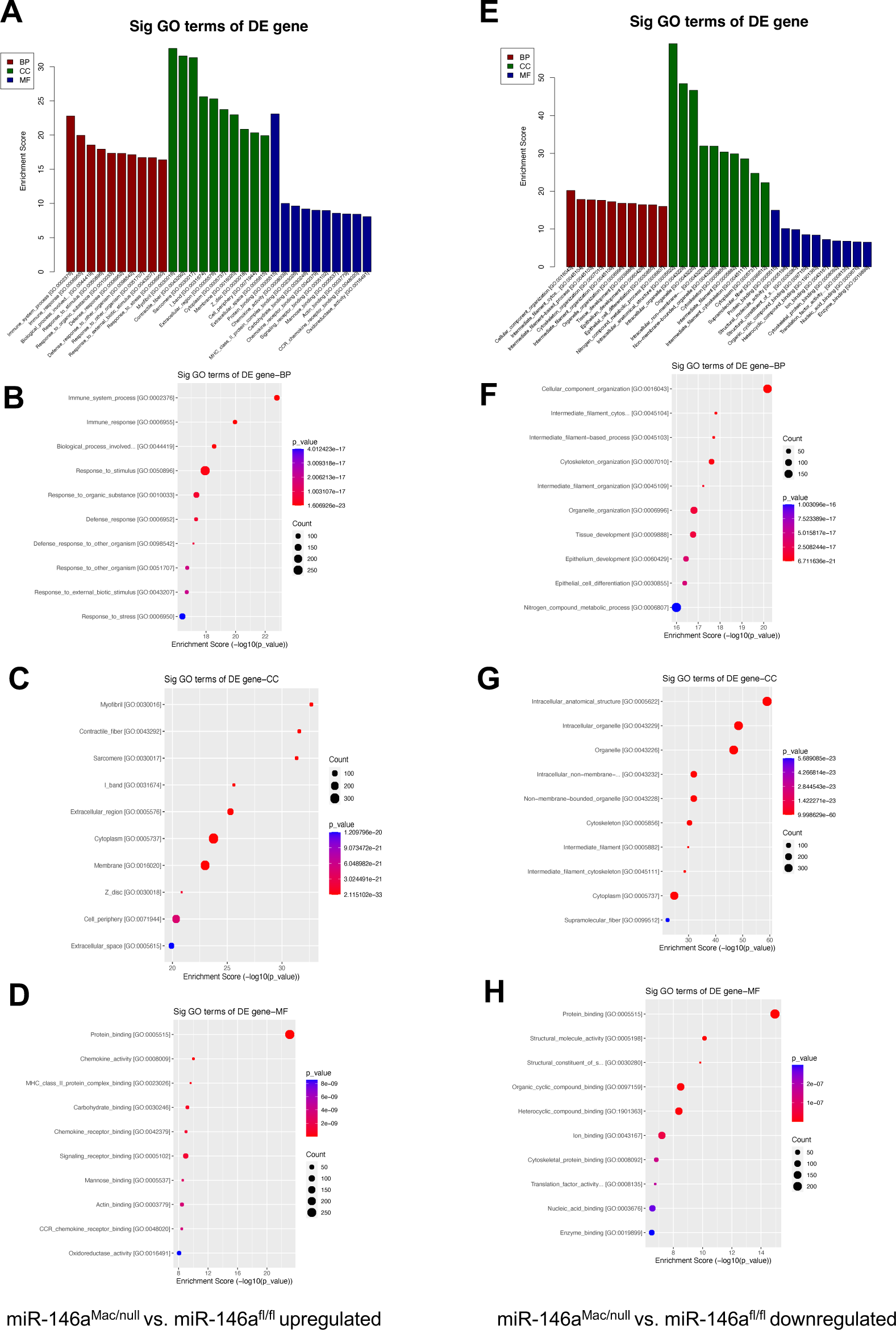
Gene Ontology (GO) enrichment analysis. (**A, E)** shows the top 10 significant GO terms in biological process (BP), molecular function (MF) and cellular component (CC) with the enrichment score in x-axis. (**B-D, F-H**) Bubble plots of the top 10 significant GO terms in BP, MF and CC displays the number of genes associated with each GO term, and the p-value color gradient scale indicates the significance of the GO term.

**Supplementary Fig. 4.**
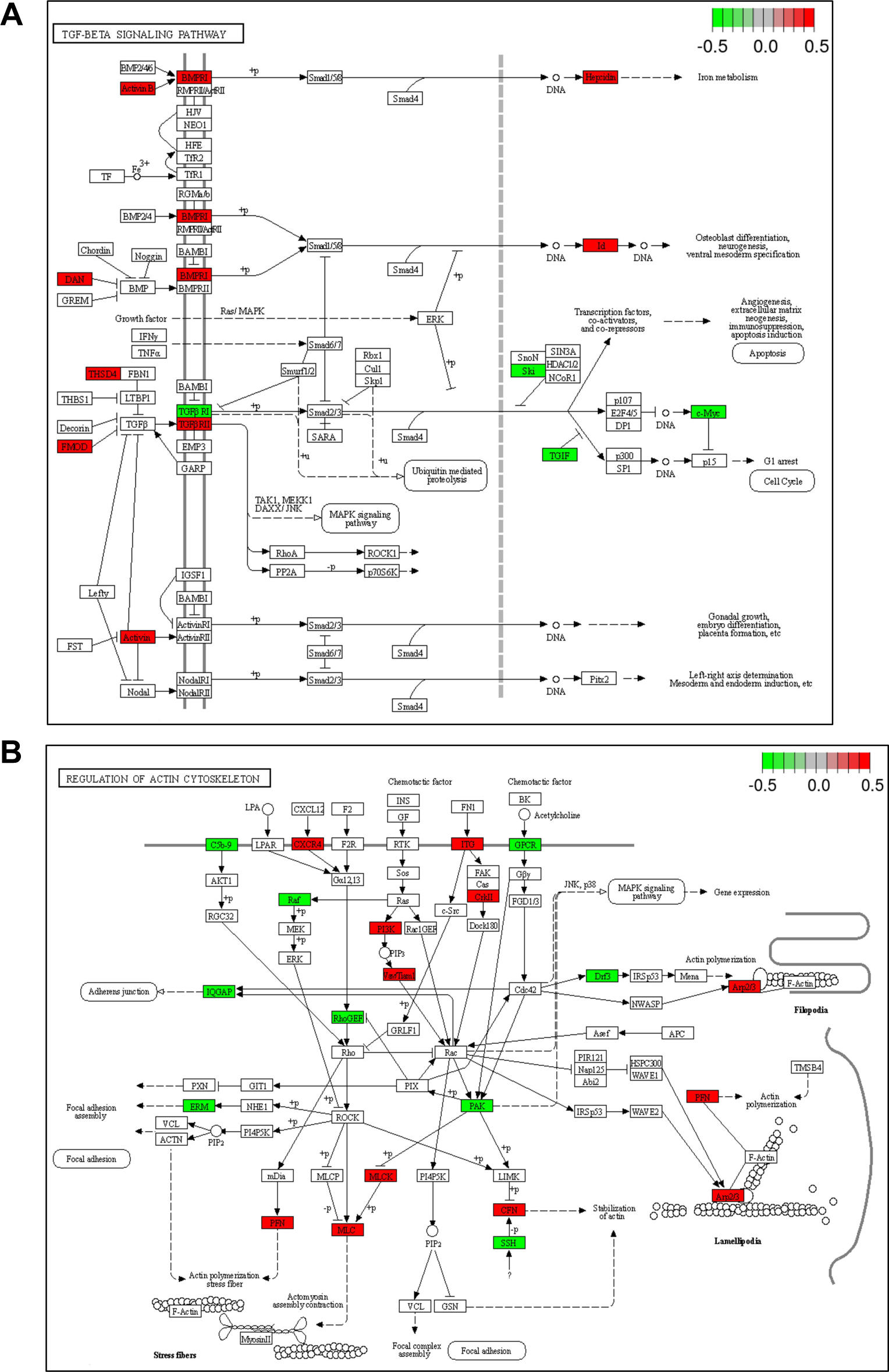
KEGG pathway maps showing dynamic interaction between the DETs. (**A**) TGFβ signaling pathway and (**B**) regulation of actin cytoskeleton is shown. The color gradient scale shows the gene expression value, bright red means upregulated, and bright green means downregulated in miR-146a^Mac/null^ as compared to miR-146a^fl/fl^.

